# Gas5A, a putative glucanosyltransferase from *Botrytis*, functions as cell death inducing protein in plants

**DOI:** 10.64898/2025.12.24.696331

**Authors:** Tobias Müller, Ida Faust, Gabriela Salinas, Charlene Chaudy, Marat Magomedov, Matthias Hahn, David Scheuring

## Abstract

The necrotrophic fungus *Botrytis cinerea*, releases numerous phytotoxic, cell death inducing proteins (CDIPs) during infection. The precise role of these proteins and their molecular function, however, is still unknown. Here, we report on the identification of a previously unknown CDIP, the glucanosyltransferase Gas5A. Functional characterization revealed that the C-terminal 60 aa of Gas5A are sufficient to induce cell death, independent from its putative enzymatic function. Gas5A localization and functional dependence on the receptor-associated kinase suppressor of BIR1-1 (SOBIR1) and the plant defense regulator ENHANCED DISEASE SUSCEPTIBILITY 1 (EDS1) indicate recognition as a pathogen-associated molecular pattern (PAMP) at the plant plasma membrane, but it is toxic also when delivered inside plant cells. Generation of a CRISPR/Cas9-assisted *Botrytis* knockout strain did not indicate any impact of Gas5A on virulence. Taken together, Gas5A represents a novel PAMP-like CDIP with additional intracellular phytotoxic activity.

## Introduction

Every year *Botrytis cinerea* (*Botrytis* hereafter), causing grey mold rot on plants, causes worldwide significant crop losses. The fungal pathogen has a broad host range and infects over 1000 different plant species from vegetables over fruits to flowers (Elad et al., 2015). Due to its socio-economic impact *Botrytis* belongs to the most studied necrotrophic fungi. The invasive life cycle is characterized by rapid killing of the host plant cells followed by colonialization of dead tissue and eventually proliferation via conidiospores (Bi et al., 2022). For the initial killing, different modes of action are employed by *Botrytis*. Upon germination on plant tissue, appressoria are formed to penetrate plant cells mechanically. Only recently, surprisingly high turgor pressures have been estimated and a unique penetration pattern for *Botrytis* was described (Müller et al., 2024). During invasion, numerous proteins, toxic metabolites and enzymes are secreted which contribute to host cell killing and decomposition (Espino et al., 2010; González-Fernández et al., 2015).

In the *Botrytis* secretome, many proteins have been characterized and classified by their function. A major group are cell wall-degrading enzymes (CWDEs), such as endopolygalacturonases (PG1-PG6) possessing pectin degrading activity (Have et al., 1998; Kars et al., 2005). Another functional class of secreted proteins are cell death inducing proteins (CDIPs). CDIPs often function by (over) activation of plant defense for the fungus’ benefit (Li et al., 2020). This includes the recognition of conserved microbial signatures, so-called pathogen-associated molecular patterns (PAMPs), by receptors at the plant plasma membrane (Boutrot and Zipfel, 2017). Binding of PAMPs to these pattern recognition receptors (PRRs) leads to the activation of PAMP triggered immunity (PTI) which can ultimately cumulate in the hypersensitive response (HR) including programmed cell death of infected tissue (Boutrot and Zipfel, 2017). Albeit the HR usually protects plants from pathogen proliferation, for necrotrophic fungi it has been shown to be beneficial (Govrin and Levine, 2000). Several CDIPs triggering the PTI and eventually leading to plant cell death has been identified in *Botrytis*. Among them are many peptides such as a 25 aa fragment from the xylanase Xyn11a (Noda et al., 2010) but also entire protein folds as from the recently found hypersensitive response inducing protein 1 (Jeblick et al., 2023). Notably, several secreted *Botrytis* enzymes are recognized as PAMPs in addition to their original function. For the aforementioned polygalacturonases as well as the xylanases Xyn11A and Xyl1, sequence-specific PAMPs have been identified and HR-like cell death was induced even if enzymatic function was impaired (Noda et al., 2010; Frías et al., 2014; Zhang et al., 2021). Notably, it has been recently reported that the secreted *Botrytis* protein Crh1 is translocated from the apoplast into plant cells acting as effector (Bi et al., 2021). However, in contrast to canonical effector functions, Crh1 induces plant defense responses and ultimately cell death.

Individual gene knockouts of secreted *Botrytis* proteins usually do not or only mildly affect the virulence, and multifold gene deletions were necessary to significantly reduce it (Leisen et al., 2022). The efficient use of CRISPR-Cas in *Botrytis* (Leisen et al., 2020) allowed to establish mutants lacking most of the described CDIPs but were still considerably aggressive on different plant species (Leisen et al., 2022). This led us to hypothesize that additional toxic proteins might exist within the secretome. During a recent secretome screening, Hip1 was identified as a CDIP and subsequently characterized (Jeblick et al., 2023). Among the other screened candidates, Bcin14g03970 also displayed toxic activity although weaker than Hip1. This gene encodes for a predicted glucanosyltransferase homologous to Gas5p of yeast which possess a signal peptide for secretion. In the present study, we report on the function of *Botrytis* Gas5A as a previously unknown CDIP. We show that the necrosis-inducing activity of Gas5A is observed when it is expressed either in the apoplast or inside plant cells. Furthermore, we demonstrate that Gas5A toxicity does not rely on its predicted function as glucanosyltransferase, but on the presence of the receptor-associated kinase suppressor of BIR1-1 (SOBIR1), indicating its recognition by a cell surface receptor. We show that Gas5A gene deletion did not result in decreased virulence on different host plants. Finaly, we hypothesize that the cell-death inducing activity was acquired as secondary function of GAS5A during plant-pathogen coevolution.

## Results

In a screen for the identification of as yet uncharacterized phytotoxic proteins, a predicted glucanosyltransferase, encoded by Bcin14g03970, was found (Jeblick et al., 2023). Using transient expression, toxicity was inconspicuous 3 days post infiltration (dpi) but significant and reproducible phytotoxic effects could be observed 4-5 dpi (Fig. 1A and B). Bcin14g03970has the highest homology to *Saccharomyces cerevisiae* 1,3-beta-glucanosyltransferase 5 (Gas5), a member of the transglycosidase GH72 family (Fig. S1). Gas proteins are conserved in most fungi, they occur as gene family consisting of 5-6 members and they function in cell-wall remodeling during vegetative growth (Rolli et al., 2011; Ao and Free, 2017). Since Gas5 has been assigned to the closely related Bcin15g05080 already (Blandenet et al., 2022), we termed Bcin14g03970 Gas5A. You could show an alignment here.

**Fig. 1.**
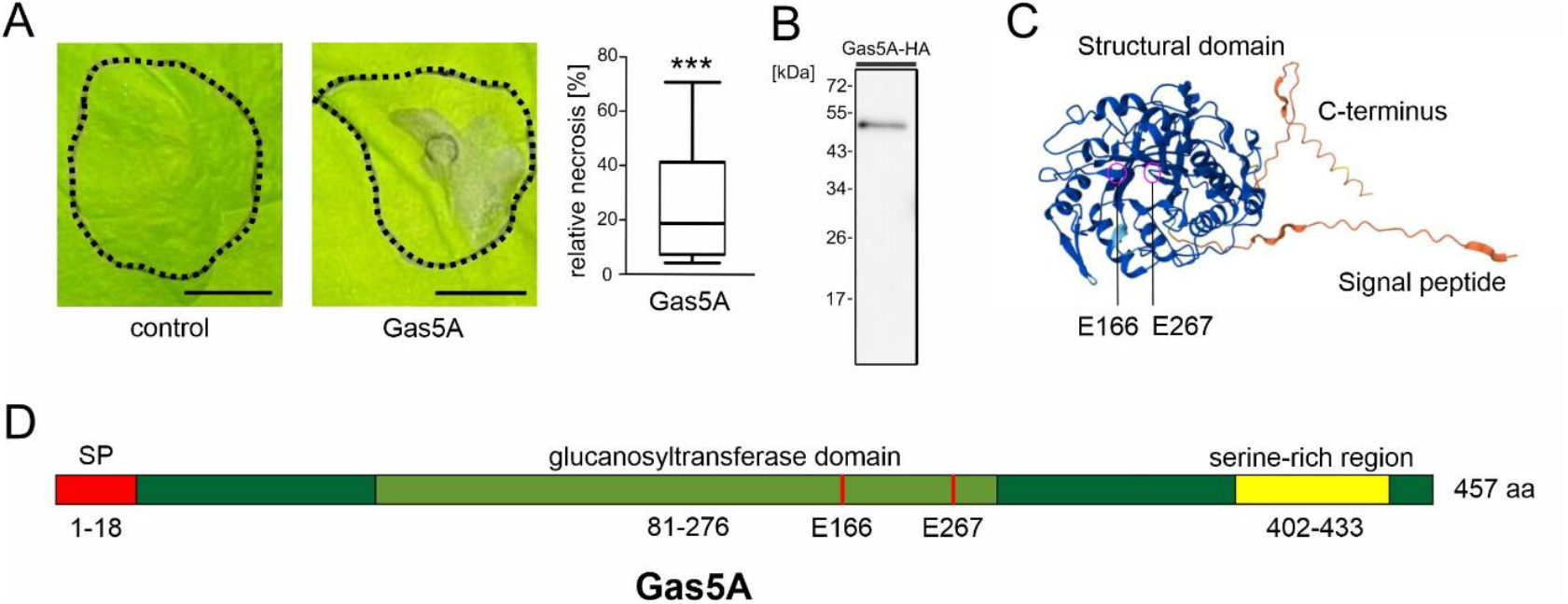
Phytotoxic activity of the putative glucanosyltransferase Gas5A. **A)** Transient expression of Gas5A in *N. benthamiana* 5 dpi. n control = 8, n Gas5A = 16. Significant differences compared with the control are shown (student t-test) followed by Tukey’s post hoc test; ^***^*P* < 0.001. Box limits in the graphs represent 25th–75th percentile, the horizontal line the median and whiskers minimum to maximum values. Scale bars= 10 mm. **B)** Westernblot confirming the expression of Gas5A. **C)** AlphaFold prediction of Gas5A protein folding. **D)** Schematic view of Gas5A.

AlphaFold predicts for Gas5A a conserved beta-barrel core forming a pore-like structure with alpha-helices surrounding the core, and a weakly conserved C-terminus extending from the core (Fig. 1C). Alignment with the Gas1 glucanosyltransferase from *S. cerevisiae* reveals two possibly conserved active sites at position E166 and E267 of *Botrytis* Gas5A (Fig. S2) (Carotti et al., 2004). This is in line with the prediction that the glucanosyltransferase activity is located between aa 81 and 276 (Fig. 1D). Notably, the C-terminus contains several serine residues and has been described as serine-rich region for yeast Gas1 (Popolo and Vai, 1999).

To assess how Gas5A mediates its phytotoxicity, we aimed to express it without signal peptide (nospGas5A) and without enzymatic activity. To this end, we replaced the glutamic acids of the active center (position E166 and E267) with structurally similar glutamines, resulting in a mutated Gas5A version (Gas5A EE). To stabilize transient expression, we co-infiltrated a binary vector coding for the viral p19 protein which suppresses plant gene silencing (Voinnet et al., 2003). In comparison to expression of Gas5A without p19 being present, we could indeed observe stronger toxicity already starting 3 dpi (Fig. 2A) Expression of Gas5A and Gas5 EE both induced similar necrosis as wildtype Gas5A, suggesting that enzymatic activity of GAS5A is not required for its phytotoxic activity (Fig. 2B).

**Fig. 2.**
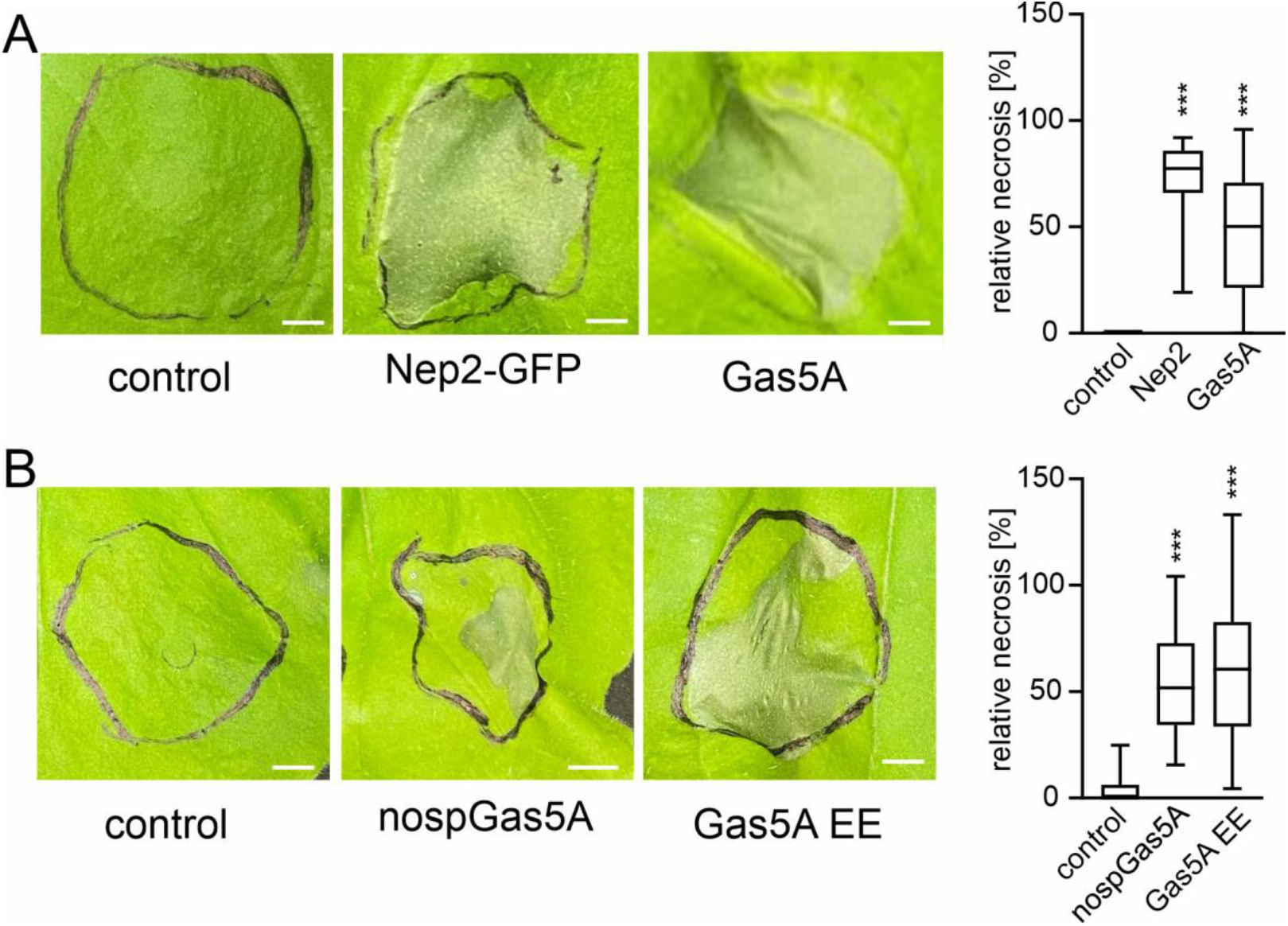
Toxicity of Gas5A is independent of its secretion and enzymatic activity. **A)** Transient expression in *N. benthamiana* and quantification of toxicity: n control = 9, n Nep2 (positive control) = 26, n Gas5A = 37. **B)** Transient expression in *N. benthamiana* and quantification of toxicity: n control = 6, n nospGas5A and Gas5A EE = 39. Significant differences compared with the control are shown (one-way analysis of variance (ANOVA) followed by Tukey’s post hoc test; ^***^*P* < 0.001. Box limits in the graphs represent 25th–75th percentile, the horizontal line the median and whiskers minimum to maximum values. Scale bars= 5 mm.

Since Gas5A was phytotoxic independently of its signal peptide, we investigated its subcellular location with and without signal peptide using GFP-fusions. Confocal microscopy of transiently expressed Gas5A-GFP and nospGas5A-GFP displayed signals at the cell borders of epidermal cells of *N. benthamiana* (Fig. 3A). To differentiate between apoplastic and cytosolic signals we used 1M NaCl to mediate plasmolysis. As the protoplast retreated from the cell wall under these conditions, fluorescence remained at the cell borders in case of Gas5A-GFP but not when nospGas5A-GFP was expressed (Fig. 3B). Colocalization with the plasma membrane marker RFP-TMD23 revealed that Gas5A-GFP is localized in close proximity (Fig. 3C). Collection of the apoplastic fluid confirmed that Gas5A-GFP is secreted into the apoplast (Fig. 3D). Notably, the addition of GFP at the C-terminus decreased Gas5A toxicity significantly (Fig. S3).

**Fig. 3.**
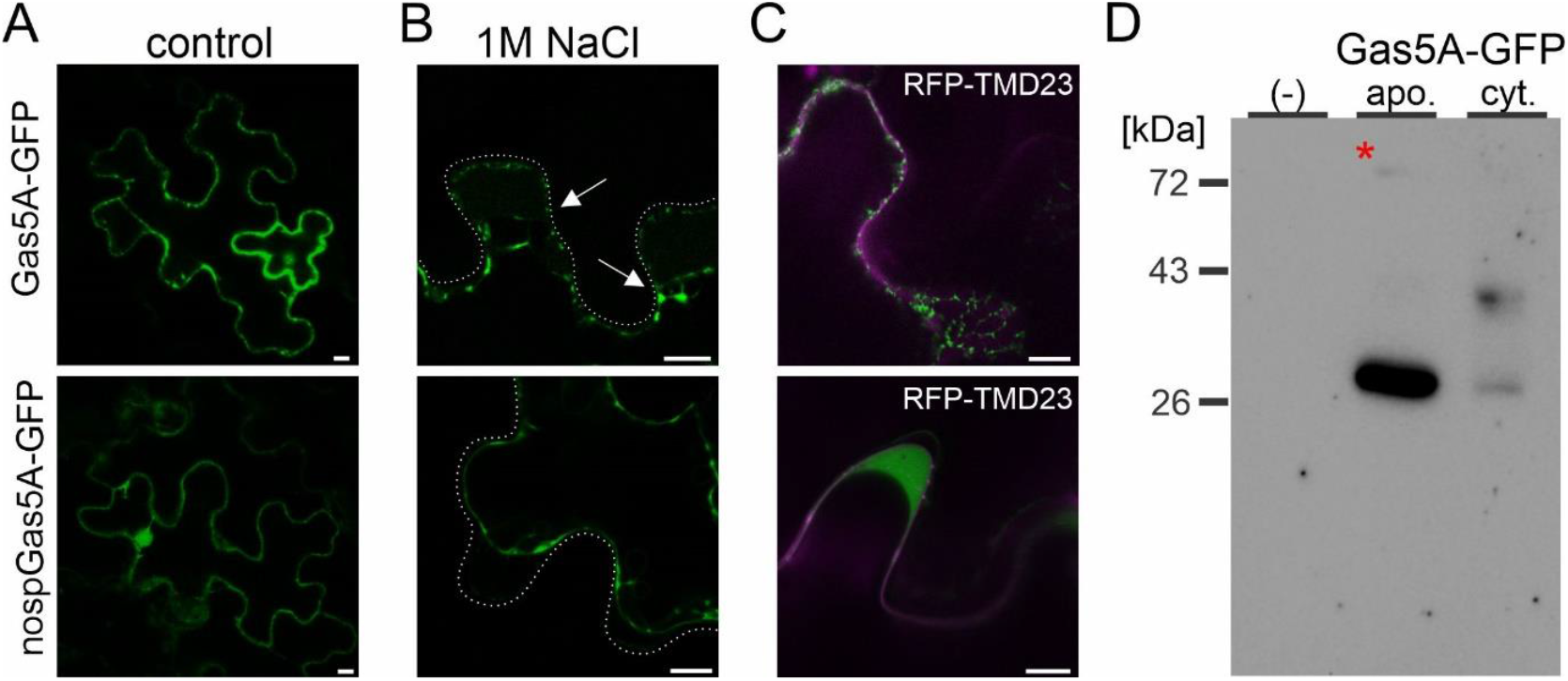
Gas5A with signal peptide (SP) is secreted. **A)** Gas5A-GFP and nospGas5A-GFP highlight the borders of epidermal *N. benthamiana* cells when transiently expressed. **B)** Induced plasmolysis by treatment with 1M NaCl. Scale bars: 10 µm. **C)** Co-expression with the plasma membrane marker RFP-TMD23. **D)** Detection of Gas5A-GFP in the apoplastic fluid. The asterisk indicates expected band size for Gas5A-GFP.

To confirm phytotoxicity observed in transient expression, we purified recombinant Gas5A using *E. coli* strain Rosetta as expression system. Despite several modifications, we were unable to express the protein in soluble form but only obtained inclusion bodies. For solubilization and refolding of the protein, we used a combination of urea and stepwise dialysis (Fig. 4A and B). Infiltration of ∼10 µM Gas5A caused necrosis in *N. benthamiana* leaves, confirming the toxic effect that was observed in transient expression (Fig. 4C).

**Fig. 4.**
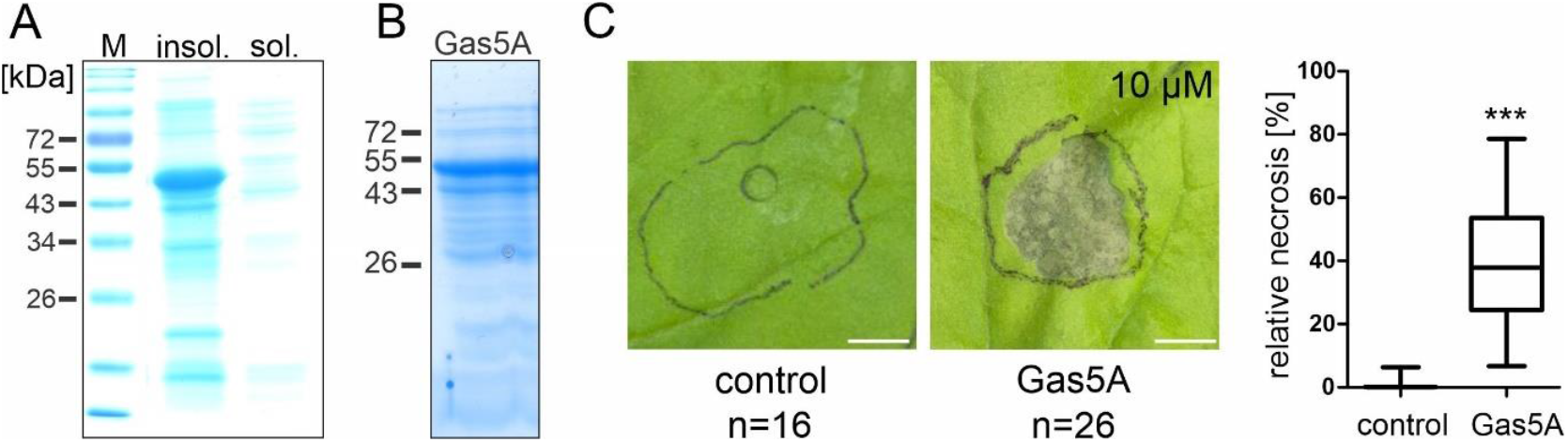
Gas5A protein expression and confirmation of toxicity. **A)** Solubility test of recombinant Gas5A expressed in *E. coli*. **B)** Gas5A after affinity purification and solubilization. **C)** Infiltration of Gas5A protein in *N. benthamiana* leaves. Scale bars:5 mm.

As Gas5A protein was toxic when secreted from transiently expressing tobacco cells, we hypothesized that it might confer toxicity by acting as a PAMP at the plant plasma membrane. Since PAMP recognition of many microbial proteins does not rely on the full aa sequence and is independent of their native fold, we constructed shortened Gas5A derivatives. Because addition of GFP to the C-terminus diminished toxicity (Fig. S3), we focused on the C-terminal part of Gas5A (Fig. 5A). Intriguingly, we could reduce length to 60 aa and still observe toxicity (Fig. 5B-D).

**Fig. 5.**
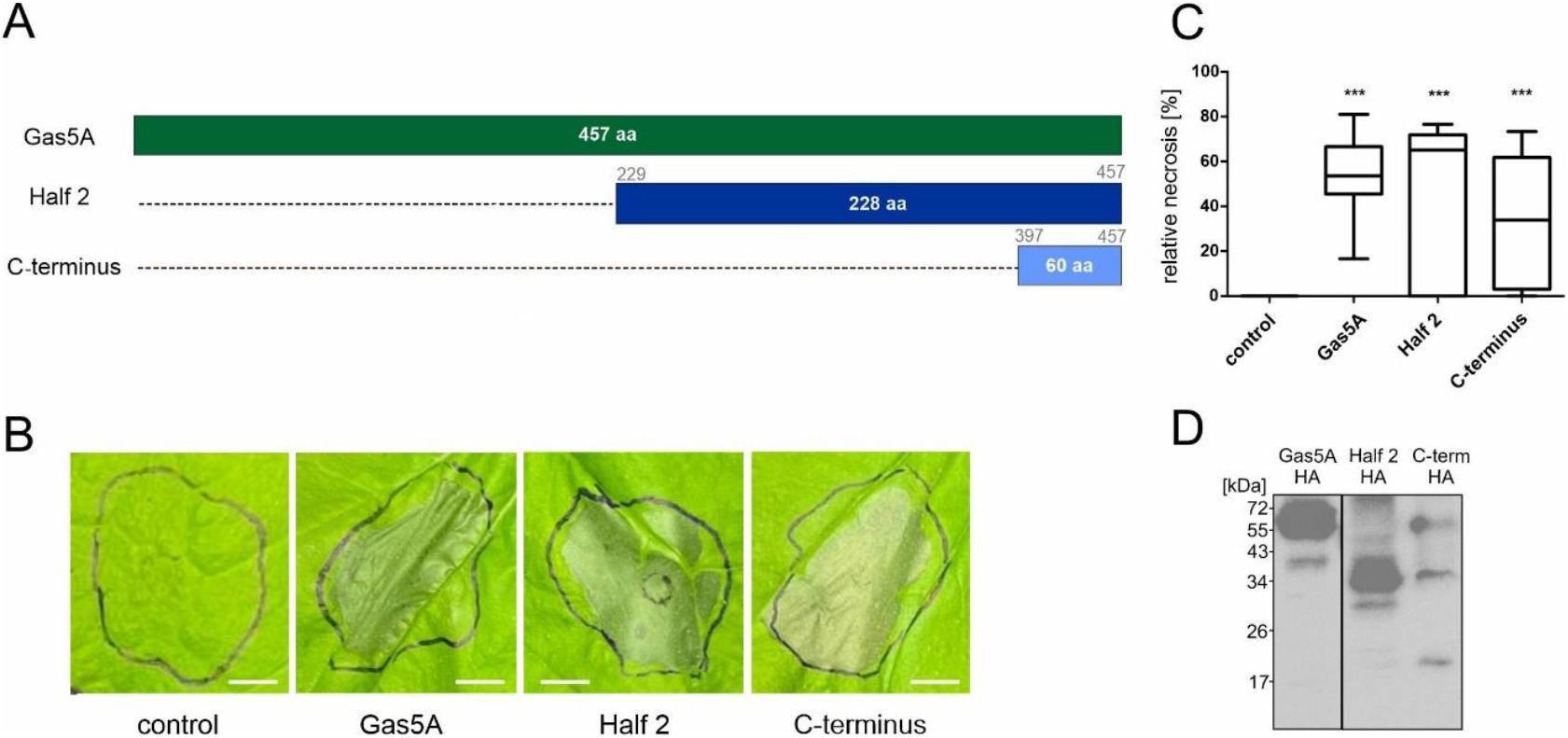
The C-terminal 60 aa of Gas5A are sufficient to confer toxicity. **A)** Overview of truncated Gas5A derivatives. **B)** Transient expression of the constructs in *N. benthamiana*. **C)** Quantification of toxicity (n=12 for each construct). Significant differences compared with the control are shown (one-way analysis of variance (ANOVA) followed by Tukey’s post hoc test; ^***^*P* < 0.001. Box limits in the graphs represent 25th–75th percentile, the horizontal line the median and whiskers minimum to maximum values. Scale bars= 5 mm. **D)** Confirmation of all constructs by immunoblotting.

To test how the plant immune system is involved to mediate Gas5A toxicity, we tested available *N. benthamiana* mutants upon transient expression. To interfere with PTI, we used a mutant of the central co-receptor suppressor of BIR1-1 (SOBIR1) (Huang et al., 2020), and to address both PTI and effector triggered immunity (ETI), we used a mutant defective in EDS1 (Lapin et al., 2019). In comparison to *N. benthamiana* wildtype plants (WT), Gas5A-mediated phytotoxicity was significantly reduced in *sobir1/sobir1-like* and *eds1* mutants while toxicity of nospGas5A was unaltered (Fig. 6A and B).

**Fig. 6.**
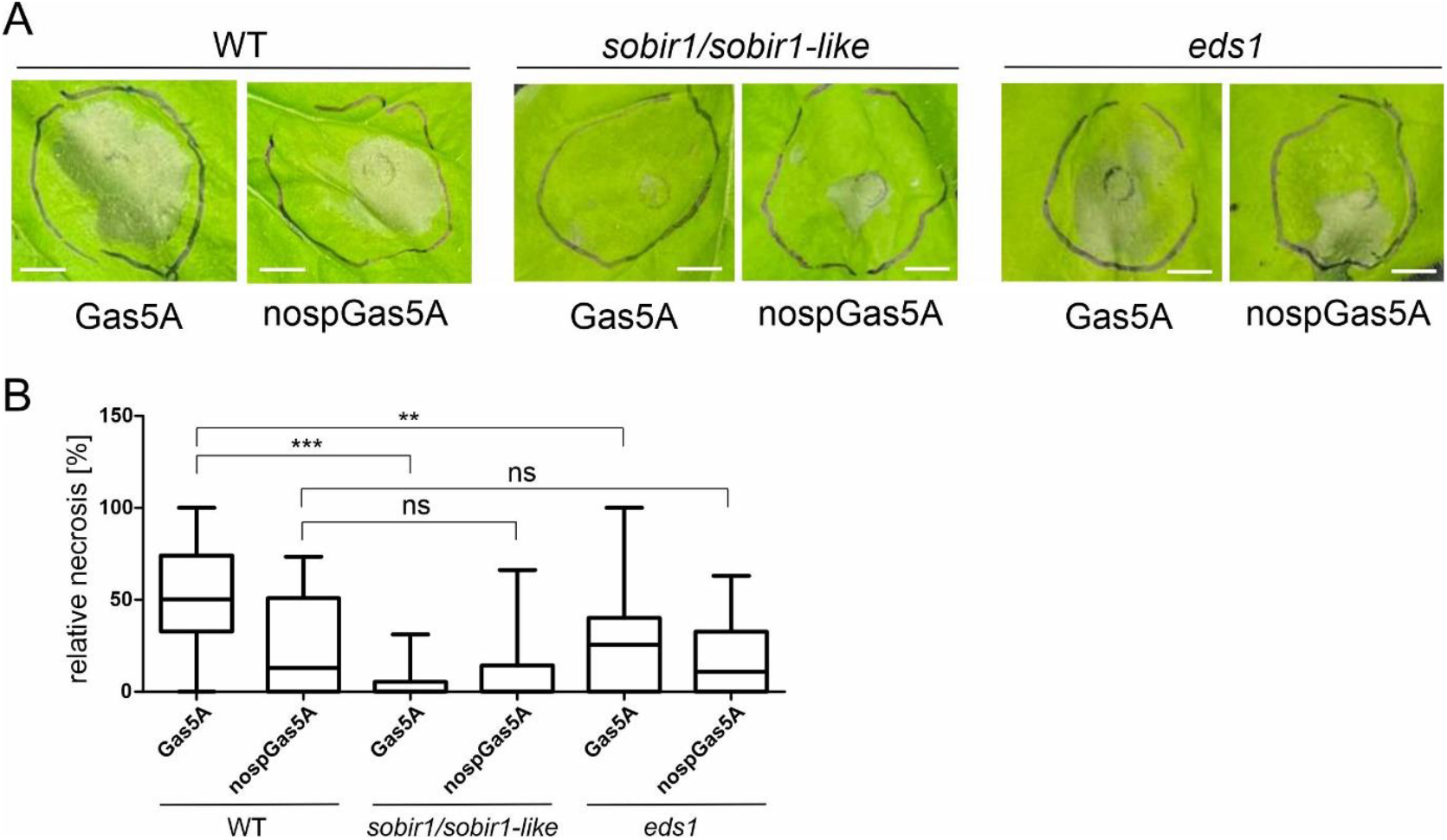
Phytotoxicity of Gas5A depends on the plant immune system. **A)** Transient Gas5A and nospGas5A expression in *N. benthamiana* WT compared to expression in *sobir1/sobir1-like* and *eds1* mutants: n Gas5A in WT = 10, n nospGas5A in WT= 6; Gas5A in *sobir1/sobir1-like* = 12, n nospGas5A in *sobir1/sobir1-like* = 12; n Gas5A in *eds1* = 10; n nospGas5A in e*ds1* = 10. **B)** Quantification: Significant differences of Gas5A and nospGas5A in WT were compared with expression in the mutants (one-way analysis of variance (ANOVA) followed by Dunnet post hoc test with WT as control; ns= not significant, ^**^*P* < 0.01. ^***^*P* < 0.001. Box limits in the graphs represent 25th–75th percentile, the horizontal line the median and whiskers minimum to maximum values. Scale bars= 5 mm.

To investigate if *Gas5A* serves as virulence factor for *Botrytis* infection, we used CRISPR/Cas9 to establish a marker-free *Gas5A* knockout strain. Growth of the *gas5A* mutant (Fig. S4) and lesion formation on leaves of tomato, *Phaseolus* bean and *Arabidopsis thaliana* were found to be unaltered when compared to the B05.10 *Botrytis* wildtype strain (Fig. 7A-C).

**Fig. 7.**
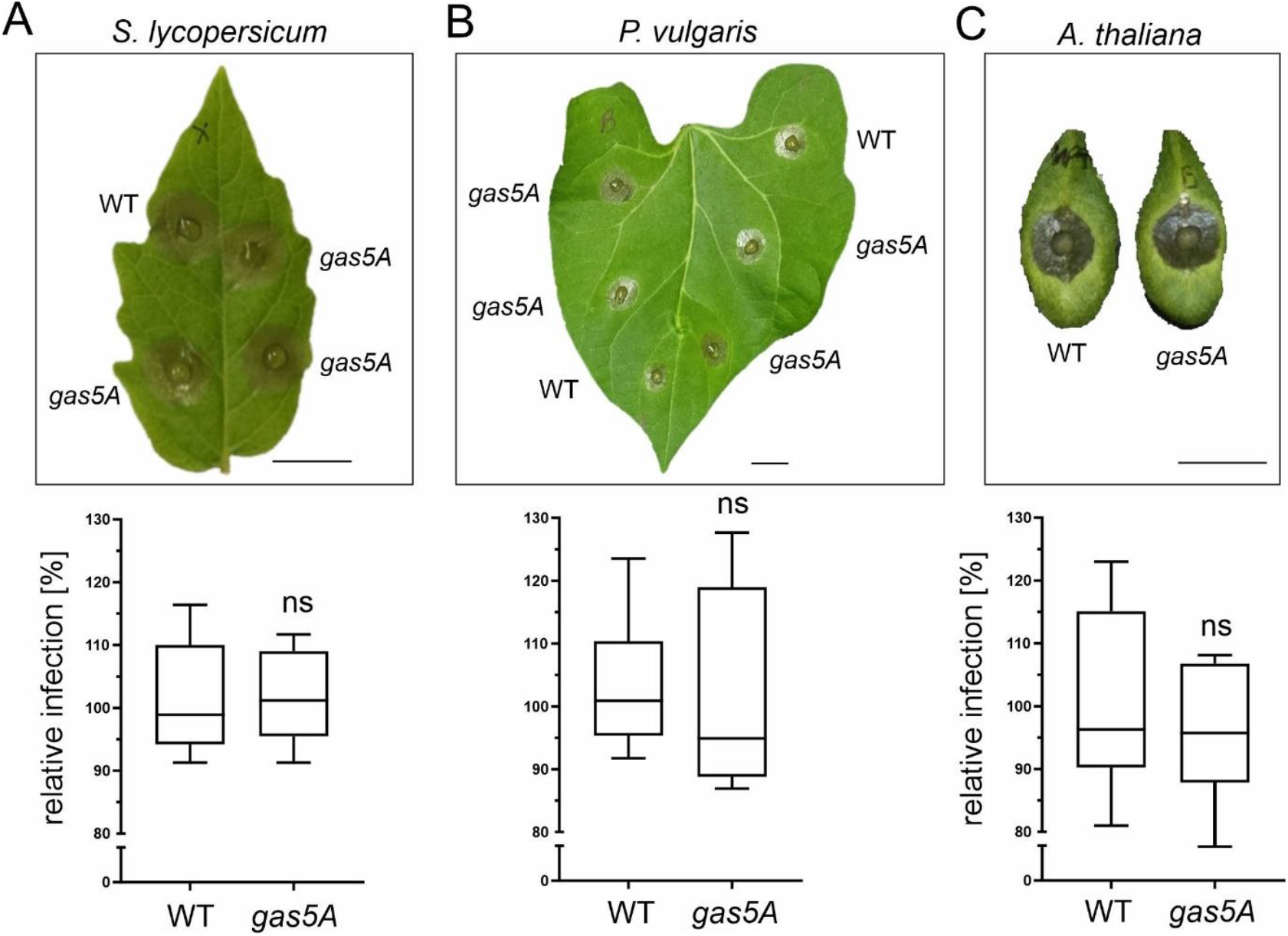
Infection test with a Gas5A knockout strain. **A)** Infection on *S. lycopersicum*: n WT = 15, n *gas5A* = 15. **B)** Infection on *P. vulgaris*: n WT = 8, n *gas5A* = 8. **C)** Infection on *A. thaliana*: n WT = 10, n *gas5A* = 15. Significant differences compared to the WT (B05.10) control are shown (unpaired t-test); ns: not significant. Box limits in the graphs represent 25th–75th percentile, the horizontal line the median and whiskers minimum to maximum values. Scale bars= 1 cm.

## Discussion

In recent years, several secreted proteins which function as cell-death inducing proteins (CDIPs) have been identified in *Botrytis*. Among them are the xylanase XYG1, the hypersensitive response-inducing protein (Hip1), the transglycosylases Crh1 and Crh4, CDI1 and Plp1 (Zhu et al., 2017; Bi et al., 2021; Jeblick et al., 2023; Zhu et al., 2023; Liang et al., 2024; Nie et al., 2025). However, knockout of single genes often did not result in decreased virulence, as reported for e.g. Hip1 and Plp1 (Jeblick et al., 2023; Liang et al., 2024). In line with this, even knockout of multiple strongly expressed genes encoding for CDIPs and toxic metabolites simultaneously did not abolish virulence completely. For a 12-fold and an 18-fold *Botrytis* mutant it was demonstrated that infection, including secondary lesion formation and sporulation, was still successful on several different plant species (Leisen et al., 2022; Müller et al., 2024). Although virulence was significantly reduced in both cases, it was concluded that secreted CDIPs function redundantly to allow for efficient infection of a broad host range (Leisen et al., 2022). This indicates that there are still unknown CDIPs secreted by *Botrytis* which might not contribute to virulence individually but to robust infection across plant species.

The finding that most *Botrytis* CDIPs are recognized by the plant immune system as PAMPs but in addition own specific functions (e.g. enzymatic activity) could suggest that their cell-death inducing activity was only acquired as secondary function during co-evolution with plants. This might be especially true for glucanosyltransferases whose primary function seems to be the modification of fungal cell walls during vegetative growth. Gas proteins are a family of enzymes with β(1,3)-glucanase/transglycosidase activity, which are crucial for the incorporation and remodeling of β(1,3)-glucan into the fungal cell wall (Kollár et al., 1997; Mouyna et al., 2000; Mouyna et al., 2005). *B. cinerea* encodes six Gas-like proteins, of which Gas5 and Gas5A are most similar to Gas5p of yeast and GEL1 of *Aspergillus fumigatus* (Fig. S1 and S2). The sufficiency of the c-terminal 60 aa of Gas5A to induce necrosis in *N. benthamiana* (Fig. 5) indicates an additional PAMP activity, eventually inducing plant cell death. Despite the high similarity of Gas5A and Gas5 which share 64.4% sequence identity, their C-termini are more diverged (33.3% sequence identity of the last 60 aa), which indicates that Gas5 might not be phytotoxic. Intriguingly, many PAMPs from *Botrytis* have been reported to induce necrosis, although this is atypical for the PTI response. The prime example of a bacterial PAMP, a peptide derived from the flagellum termed flg22, is not inducing cell death but rather enhances plant resistance (Zipfel et al., 2004). For secreted *Botrytis* proteins, however, several cell death-inducing PAMPs have been reported. This includes peptides from the xylanases Xyl1 (26 aa) and Xyn11A (30 aa), IEB1 (35 aa) and Spl1 (40 aa) (Noda et al., 2010; Frías et al., 2011; Frías et al., 2016; Yang et al., 2018).

Another line of evidence for Gas5A acting as cell-death inducing PAMP is the dependence on SOBIR1 and EDS1 signaling (Fig. 6). Dependency on the LRR receptor-like kinases SOBIR1 and BAK1 had been shown previously for other CDIPs like Xyl1, XYG1 and Plp1 (Zhu et al., 2017; Yang et al., 2018; Nie et al., 2025). This suggests that Gas5A is recognized by a receptor-like protein at the cell surface requiring at least SOBIR1 as co-receptor for signaling. Interference with the formation of the Gas5A-RLP-SOBIR1 complex hence interrupts signaling and suppresses cell-death induction.

Notably, Gas5A is toxic even in the absence of a signal peptide, raising the question of whether PTI induction is its sole mechanism for triggering cell death. Recent studies indicate that some CDIPs secreted into the apoplast are subsequently translocated into plant cells, leading to cell death and activation of plant immunity, such as SSP2 (Zhu et al., 2022) and Crh1 (Bi et al., 2021). For the secreted *Botrytis* transglycosylase Crh1 two different regions of the protein are responsible for uptake and cell death induction. In another transglycosylase, Crh4, also two functional distinct domains are present, one enhancing cell death and PTI the other inducing a weaker response (Liang et al., 2024). Thus, we cannot exclude that a portion of Gas5A might be translocated into the plant’s cytosol and exerts a second cell-death inducing function there.

Gene deletion of many CDIPs in *Botrytis* does not lead to virulence penalties. Gene knockout of IEB1, XYG1, Hip1 and Plp1 e.g. did not result in reduced virulence (Frías et al., 2016; Zhu et al., 2017; Jeblick et al., 2023). This was also true for Gas5A, where infection with a deletion strain on several host plants did not decrease infection rate (Fig. 7). This suggests that the (relatively low) phytotoxic activity of Gas5A does not significantly contribute to the overall phytotoxic activity and virulence of *Botrytis*. Furthermore, the normal growth of the *gas5A* mutant indicates that Gas5A does not fulfil an essential function, or that its loss can be compensated by its homologue Gas5 or other Gas proteins. This might be explained by the high redundancy of phytotoxic compounds secreted by *Botrytis* as experiments with higher-order mutants imply (Leisen et al., 2022). Nevertheless, the toxic activity of Gas5A might contribute to robust infection of a wide range of different plant species, even under suboptimal conditions.

## Materials & methods

### Plant Material and Growth Conditions

For transient expression and leaf infiltration *N. benthamiana* was grown for 4–6 weeks on soil at 23°C under long-day conditions (14-h light/10-h dark). For infection tests, *P. vulgaris* (genotype N9059), *N. benthamiana*, and tomato (*S. lycopersicum*) ‘Marmande’ were grown under similar conditions, while Arabidopsis (*A. thaliana*) ecotype Col-0 was grown under short-day regime (8-h light/16-h dark) for 5 weeks at 22°C.

### Cultivation of *Botrytis* and infection tests

*Botrytis cinerea* B05.10 was used as wild-type strain for infection tests, growth tests and as genetic background for *gas5A* knockout. Cultivation of *Botrytis* was performed as described previously (Müller *et al*., 2018). Infection tests were carried out using 1 × 10^5^ spores mL^−1^ for tomato and *P. vulgaris* and 1 × 10^5^ spores mL^−1^ for Arabidopsis. For infection of Arabidopsis, spores were pregerminated for 6 h in semi-solid medium (0.35% agar). In general, lesions were photographed after 48 h and lesion size was quantified (area) using the freehand tool in ImageJ.

### Recombinant plasmid construction

Coding sequences were amplified from *Botrytis* cDNA synthesized from total RNA extracts. Full-length as well as truncated versions of Gas5A were generated by a Phusion PCR protocol using the corresponding primer pairs. Destination vector and PCR fragments were digested with the Type IIS restriction enzyme Bsa1 at the indicated sites (Supplementary Table) and ligation reactions were performed using T4 DNA ligase (New England BioLabs). For transient expression, a GreenGate cloning system was employed (Lampropoulos *et al*., 2013), pGGZ003 was used as destination vector, pGGZ003 harbors a cauliflower mosaic virus (CMV) 35S promoter, an N-terminal signal peptide (SP), an RCBS terminator, an HA-tag for immunological detection, and a spectinomycin resistance cassette. To generate the Gas5A construct lacking a signal peptide (nospGas5A), the destination vector without the N-terminal SP module was used and a start codon introduced upstream of the coding sequence. The Gas5A–GFP fusion construct was assembled by GreenGate cloning by adding a GFP tag at the C-terminus using the reverse primers.For recombinant protein expression, the Gas5A coding sequence lacking the signal peptide was amplified and cloned into the pET28a (+) expression vector, containing an N-terminal His-tag fusion as well. All gene constructs were verified by Sanger sequencing (Seq-It, Germany). All primers used are shown in the table below. Gene constructs were verified by sequencing (Eurofins Genomics, Light Run, Germany).

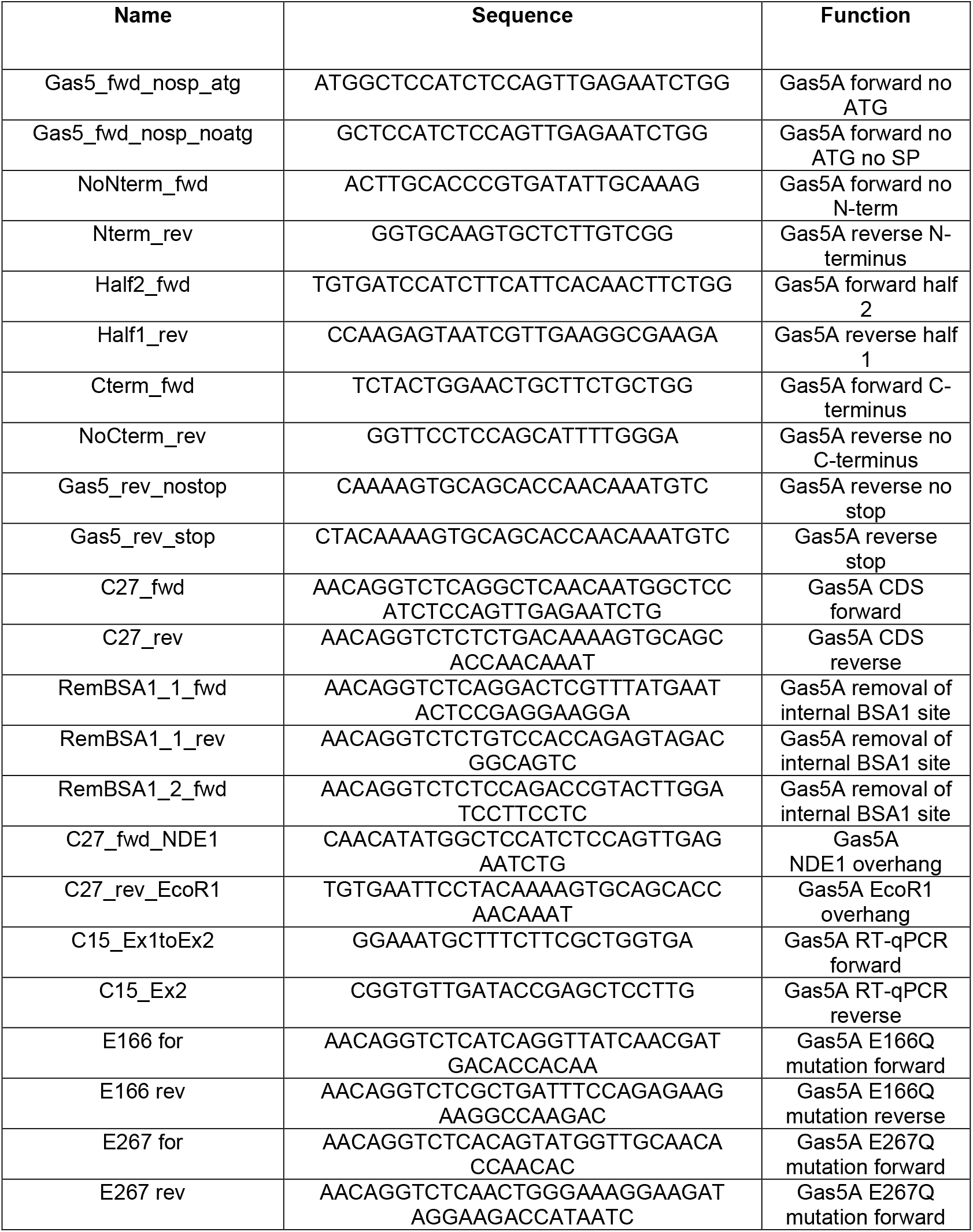

### CRISPR/Cas9-mediated generation of *Botrytis* knockout strains

CRISPR/Cas9-mediated gene deletion in Botrytis cinerea was performed using ribonucleoprotein (RNP) complexes as described previously with minor modifications (Leisen *et al*., 2020). For targeted gene knockout, two single guide RNAs (sgRNAs) were designed to introduce double-strand breaks upstream and downstream of the coding region, resulting in complete deletion of the target gene. sgRNAs were synthesized using the following oligonucleotides: AAGCTAATACGACTCACTATAGGAGATAACATATGGATAGTGGGTTTTAGAGCTAGAAATAGCAAG (NS10-gltr1gRNA-L1) and AAGCTAATACGACTCACTATAGGAGGCATGGAGAGAGGCTGCGGTTTTAGAGCTAGAAATAGCAAG (NS13-gltr1gRNA-R1). Transformation of *Botrytis* was performed as described previously (Leisen *et al*., 2020). For PCR-based verification of *Gas5A* knockout, primers located outside and inside the coding sequence (Fig. S5) were used:

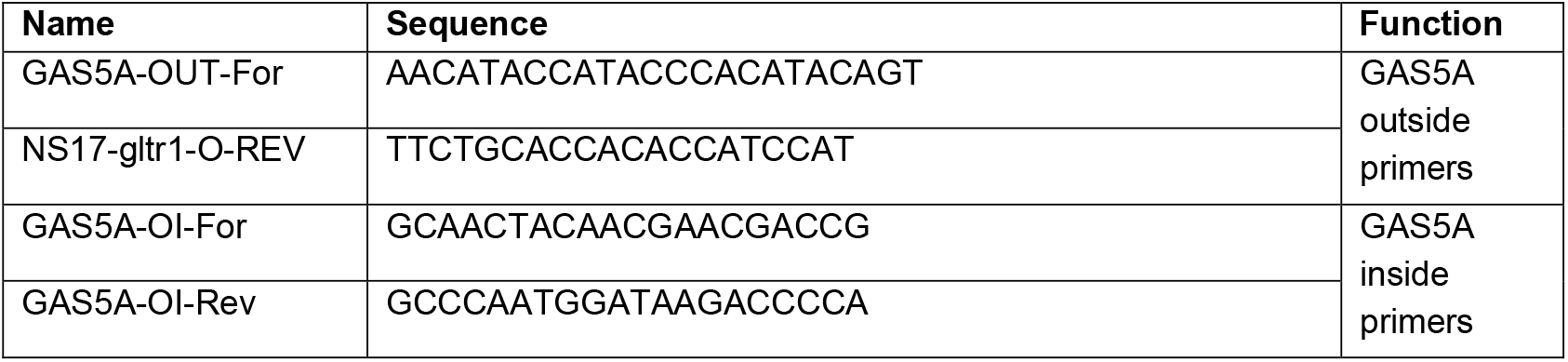

### *In silico* motif analysis and structural modeling

The amino acid sequence of Gas5A was analyzed using MOTIFinder (https://www.genome.jp/tools/motif/) to predict potential enzymatic activity. This computational tool aligns sequences and scans them for functional motifs, enabling the identification of regions associated with biological functions. To further investigate the structural context of Gas5A, modelling was performed using AlphaFold. The obtained predicted structural model allowed the visualization of the spatial organization of the protein as well as the model confidence using the predicted Local Distance Difference scores. Signal peptides were determined with SignalP 5.0 (https://services.healthtech.dtu.dk/services/SignalP-5.0/).

### Transient gene expression

For transient expression, *A. tumefaciens* (strain GV3101::mp90) was electroporated in the presence of 100 ng vector using the gene pulser II (Bio-Rad, USA) set to 2.5 kV, 25 µF, and 400 Ω. Overnight grown liquid cultures of transformed agrobacteria were centrifuged at 2800 rpm for 10 minutes and washed twice. The final supernatant was discarded, and the pellet resuspended in 2 ml infiltration buffer containing 250 mg D-glucose, 5 ml 20 mM sodium phosphate, 5 ml 0,5 M 2-Morpholinoethanesulphonic acid, 50 μl acetosyringone and up to 50 ml water. The optical density (OD) was adjusted to 0.3 prior infiltrations in *N. benthamiana*. Before infiltration, the suspensions were mixed with a p19-suspension in a 1:1 ratio. Subsequently, the infiltrated area was marked and assessed after 96 h incubation. Relative necrotic area was calculated as ratio of the infiltrated area and the necrotic area. For co-expressions, agrobacteria o\n cultures were adjusted to OD 0.15 and mixed before infiltration.

### Protein extraction and immunological detection

Proteins were extracted from leaf disks (20 mm diameter) of infiltrated *N. benthamiana* by grinding in liquid nitrogen using a mortar and pistil. The frozen powder was suspended in extraction buffer (100 mM potassium, 20-mM 4-(2-hydroxyethyl)-1-piperazineethanesulfonic acid (HEPES), 0.1-mM ethylenediaminetetraacetic acid (EDTA), 2-mM DTT, 1-mM phenylmethylsulfonyl fluoride (PMSF); pH 7.5) and after centrifugation protein determination by Bradford was carried out. Equal protein amounts were separated via sodium dodecylsulfate polyacrylamide gel electrophoresis (SDS–PAGE) and subsequently subjected to immunoblot on nitrocellulose membrane. Monoclonal antibody against HA coupled to peroxidase (Roche; Cat. No. 12013819001, 1:1000 dilution) or a polyclonal antibody against GFP (Roche;11814460001, 1:500 dilution) were used for chemiluminescence detection (ECL Prime Kit; Amersham, UK) on a ChemiDoc detection system (Bíorad, USA).

### Recombinant protein expression

*Escherichia coli* strain T7 shuffle (New England Biolabs, USA) was used to express Gas5A. Induction of protein expression was enabled by 0,5 mM isopropyl β-d-1-thiogalactopyranoside (IPTG) and conducted at 37°C for 3h. Solubility was checked using B-PER™ Reagent (Thermo scientific, USA). After bacterial lysis, protein purification was performed via immobilized metal affinity chromatography using His/Ni beads (Roth, Germany) as resin. To ensure solubility throughout all necessary buffers contained 6M Urea. Thereafter highspeed centrifugation was carried out to concentrate protein and remove urea using Amicon ultra centrifugal filter devices (Milipore, USA) and 50 mM phosphate buffer with 150 mM NaCl (pH 6).

### Isolation of apoplastic fluid after transient expression

72h after transformation, *N. benthamiana* leaves were incubated in 500 ml ddH_2_O while a 60 mbar vacuum was applied for 10 min. This was repeated twice and then the leaves were carefully placed in a syringe without needle and centrifuged at 2,000*g* and 4°C for 45 min to collect the apoplastic fluid. For immunological detection, the apoplastic fluid was run on an SDS-PAGE and GFP was detected after western blotting using a GFP antibody o/n (Roche;11814460001, 1:1000 dilution). As secondary antibody, a goat-anti mouse peroxidase conjugate (Calbiochem; 401215) was used in 1:5000 dilution for 1h at 4°C.

### Confocal microscopy

Images were acquired using a Zeiss LSM880 AxioObserver confocal laser scanning microscope equipped with a Zeiss C-Apochromat 40×/1.2 W AutoCorr M27 water-immersion objective (INST 248/254-1). Fluorescent signals of GFP (excitation/emission 488 nm/500–571 nm) and red fluorescent protein (RFP) (excitation/emission 594 nm/600–732 nm) were processed using the Zeiss software ZEN 2.3 or ImageJ (https://imagej.nih.gov/ij/). Laser intensity was set to 2% (594) and 3% (488) and the gains were adjusted in accordance with the protein expression level. For co-localization studies, the plasma membrane marker RFP-TMD23 (pFK44) was used (Scheuring *et al*., 2012)

## Statistical analysis

Analysis was carried out using the GraphPad Prism 9 software. The detailed statistical method employed is provided in the respective figure legends. All experiments were carried out at least three times. Box limits in the graphs represent 25th to 75th percentile, the horizontal line the median and whiskers minimum to maximum values.

## Acknowledgements

This work was supported by grants from the *BioComp* research initiative (Rhineland-Palatinate, Germany) and the German research foundation (DFG; SCHE 1836/5-1) to MH and DS.

## Contributions

TM, IF, GS, CC, MM and DS performed and analysed experiments. TM, GS, CC, MM and DS designed the figures and performed statistical analysis. MH and DS conceived the study and DS wrote the manuscript. All authors saw and commented on the manuscript.

## Competing interests

The authors declare no competing interests.

## Notes

### Competing Interest Statement

The authors have declared no competing interest.

