## Supplementary Figures for "Gas5A, a putative glucanosyltransferase from *Botrytis*, functions as cell death inducing protein in plants"

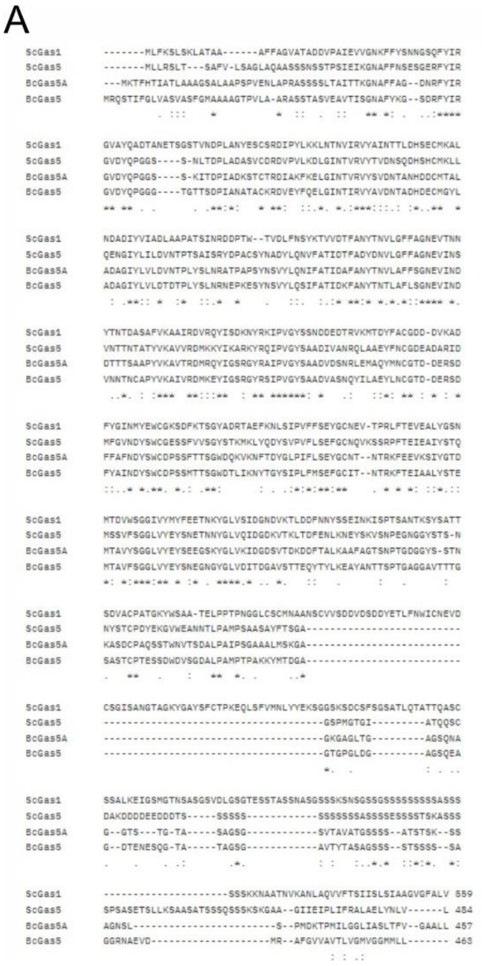

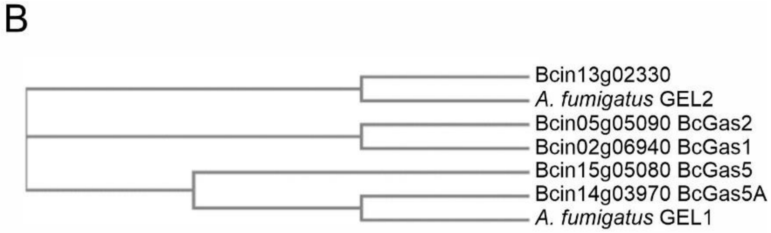


C

| Scientific Name | Max Score | Total Score | Query Cover | E value | Per. Ident | Acc. Len | Accession |
|---|---|---|---|---|---|---|---|
| BcGas5A | 924 | 924 | 100% | 0.0 | 100.00% | 457 | XP_001550496.1 |
| BcGas5 | 537 | 537 | 89% | 0.0 | 64.41% | 463 | XP_024553506.1 |
| Bcin13g02330 | 291 | 291 | 76% | 2e-92 | 43.90% | 492 | XP_024552541.1 |
| BcGas1 | 292 | 292 | 73% | 4e-92 | 45.06% | 540 | XP_001547403.1 |
| BcGas2 | 229 | 229 | 74% | 3e-69 | 39.01% | 436 | XP_024548840.1 |
| Bcin01g09460 | 154 | 154 | 55% | 5e-42 | 36.40% | 288 | XP_024546600.1 |

**Supplementary Fig. S1: Relationship of fungal Gas proteins. A)** Amino acid alignment (Clustal W) Gas1 and Gas5 from *Saccharomyces cerevisiae* with Gas5 and Gas5A from *Botrytis cinerea.* **B)** CLUSTAL O (1.2.4) multiple sequence alignment: BcGas5/ BcGas5A are orthologs of *A. fumigatus* GEL1, Bcin13g02330 is the ortholog of *A. fumigatus* GEL2. **C**) *B. cinerea* contains 6 Gas proteins.

```
Gas5A       MKTFHTIATLAAAGSALAAPSPVENLAPRASSSSLTAITTKGNAFFAG--DNRFYIRGVD 58
Gas1        -MLFKSLSKLATAAAF---------FAGVATADDVPAIEVVGNKFFYSNNGSQFYIRGVA 50
               *::::.**:*.:          :*  *::..: ** . ** ** .  ..:******

Gas5A       YQPGG-----SSKITDPIADKSTCTRDIAKFKELGINTVRVYSVDNTANHDDCMTALADA 113
Gas1        YQADTANETSGSTVNDPLANYESCSRDIPYLKKLNTNVIRVYAINTTLDHSECMKALNDA 110
            ** .      .*.:.**:*: .:*:***  :*:*. *.:***:::.* :*.:**.** **

Gas5A       GIYLVLDVNTPLYSLNRATPAPSYNSVYLQNIFATIDAFANYTNVLAFFSGNEVINDDTT 173
Gas1        DIYVIADLAAPATSINRDDPT--WTVDLFNSYKTVVDTFANYTNVLGFFAGNEVTNNYTN 168
            .**:: *: :*  *:**  *:  :.   ::.  :.:*:********.**:**** *: *.

Gas5A       TSAAPYVKAVTRDMRQYIGSRGYRAIPVGYSAADVDSNRLEMAQYMNCGTDDERSDFFAF 233
Gas1        TDASAFVKAAIRDVRQYISDKNYRKIPVGYSSNDDEDTRVKMTDYFACGDDDVKADFYGI 228
            *.*: :***. **:****..:.** ******: * :..*::*::*: ** ** ::**:.:

Gas5A       NDYSWCDPSSFTTSGWDQKVKNFTDYGLPIFLSEYGCNTN-TRKFEEVKSIYGTDMTAVY 292
Gas1        NMYEWCGKSDFKTSGYADRTAEFKNLSIPVFFSEYGCNEVTPRLFTEVEALYGSNMTDVW 288
            * *.**. *.*.***: ::. :*.: .:*:*:******    * * **:::**::** *:

Gas5A       SGGLVYEYSEEGSKYGLVKIDGDSVTDKDDFTALKAAFAGTSN-PTGDGGYSSTNKASDC 351
Gas1        SGGIVYMYFEETNKYGLVSIDGNDVKTLDDFNNYSSEINKISPTSANTKSYSATTSDVAC 348
            ***:** * ** .*****.***:.*.  ***.  .: :   *   :.  .**:*..   *

Gas5A       PAQSSTWNVTSDALPAIPSGAAALMSKGAGK-----------------------GAGLTG 388
Gas1        PATGKYWSAATELPPTPNGGLCSCMNAANSCVVSDDVDSDDYETLFNWICNEVDCSGISA 408
            ** .. *..:::  *:  .* .: *. . .                         :*::.

Gas5A       AGSQ------------------------NAGGTST------------------------- 399
Gas1        NGTAGKYGAYSFCTPKEQLSFVMNLYYEKSGGSKSDCSFSGSATLQTATTQASCSSALKE 468
             *:                         ::**:.:

Gas5A       ----------GTASAGSGSV------------TAVATGSSSSATSTSKSSAGNSLSPMDK 437
Gas1        IGSMGTNSASGSVDLGSGTESSTASSNASGSSSKSNSGSSGSSSSSSSSSASSSSSSKKN 528
                      *:.. ***:             :   :***.*::*:*.***..* *  .:

Gas5A       TPMILGGLIASLTFV-----------GAALL      457
Gas1        AATNVKANLAQVVFTSIISLSIAAGVGFALV      559
            :   : . :*.:.*.           * **:
```

**Supplementary Fig. S2:** Conserved glutamic acid residues of Gas5A (*Botrytis cinerea)* and Gas1 (*Saccharomyces cerevisiae)*. E166 and E267 are marked in red.

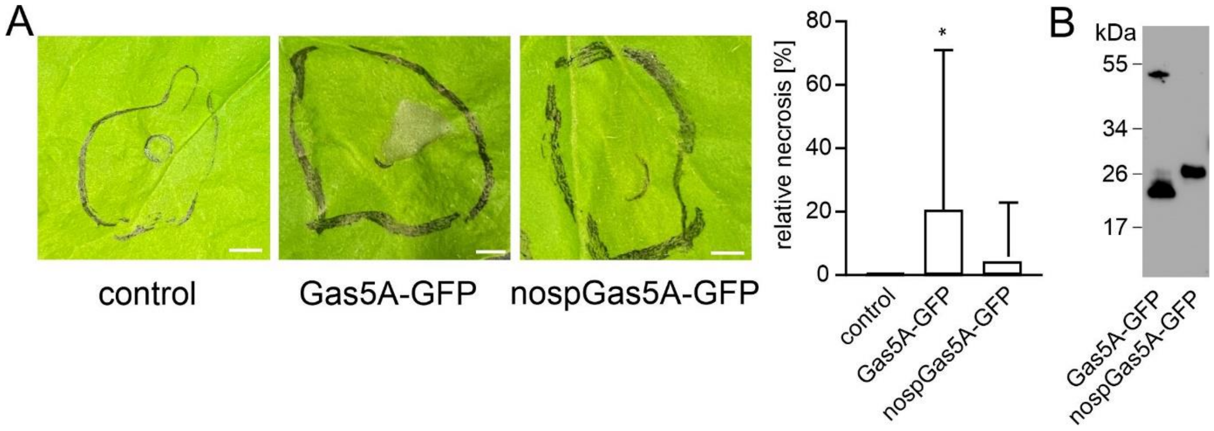


**Supplementary Fig. S3: GFP fusion to the GAS5A C-terminus attenuates toxicity. A)** n control = 9, n Gas5A-GFP: 25, n nospGas5-GFP = 29. Significant differences compared with the control are shown (one-way analysis of variance (ANOVA) followed by Tukey's post hoc test; ns: not significant. Box limits in the graphs represent 25th–75th percentile, the horizontal line the median and whiskers minimum to maximum values. Scale bars= 5mm.

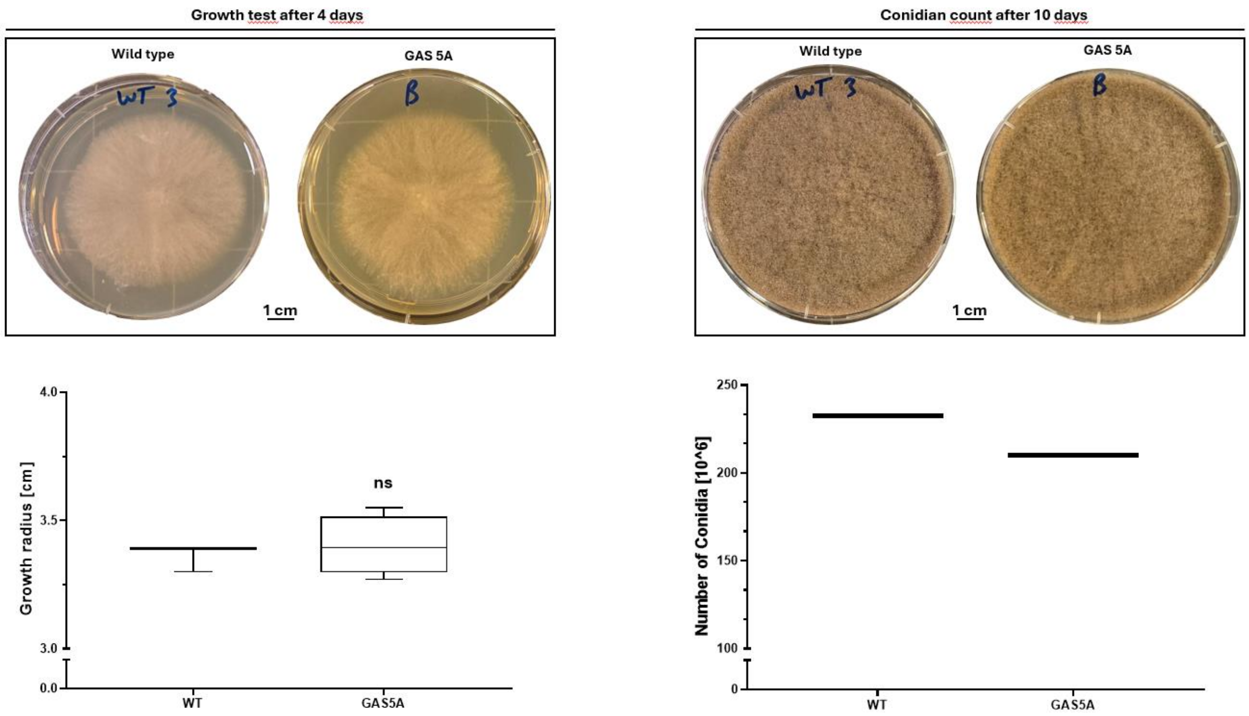


**Supplementary Fig. S4: Botrytis knockout *gas5A* strain behaves like the B05.10 wildtype (WT).**

**Single K.O.-Mutation Verification**

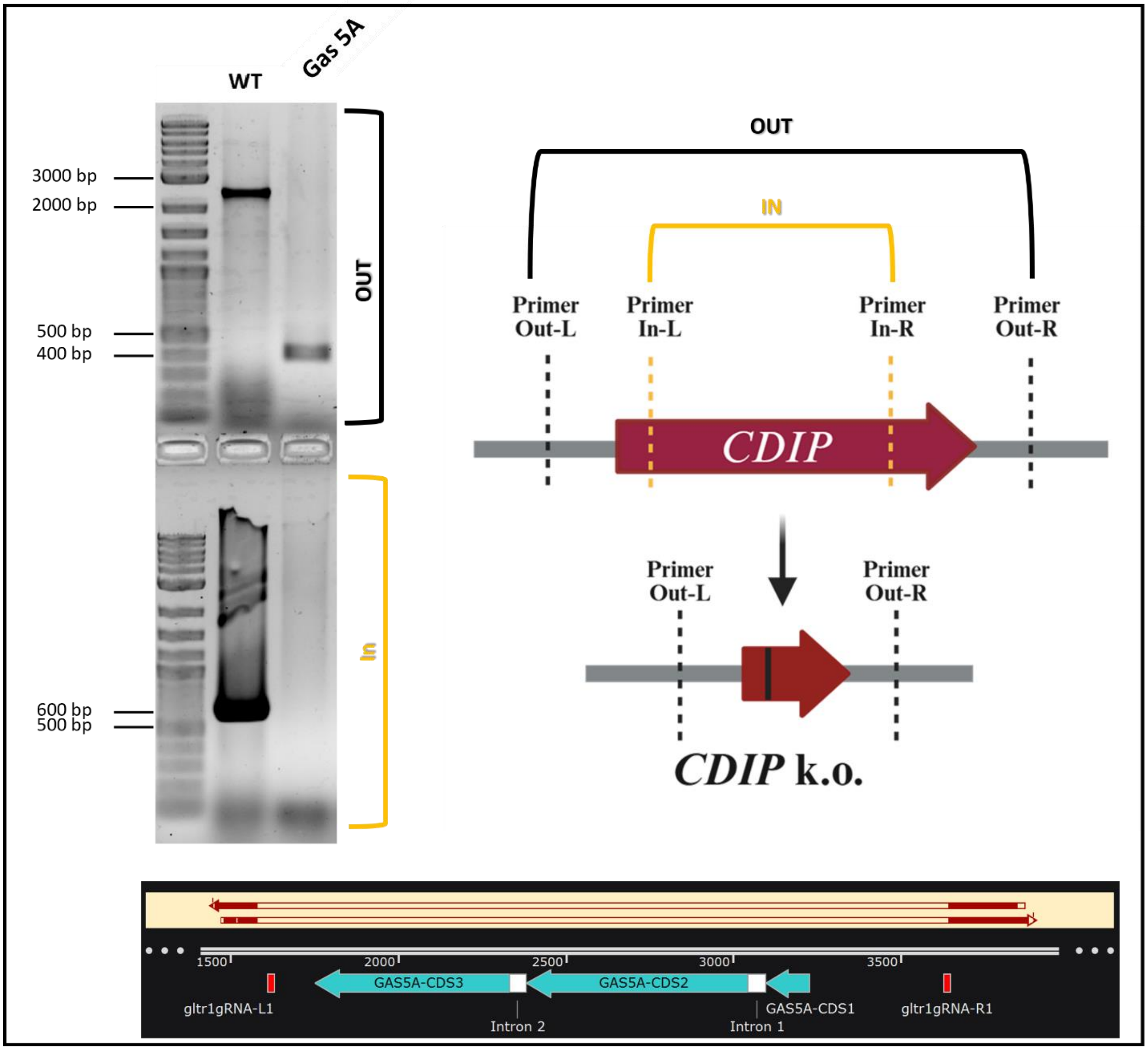


**Supplementary Fig. S5: Confirmation of *Botrytis Gas5A* knockout.**
